# A joint framework for studying population structure using principal component analysis and F-statistics

**DOI:** 10.1101/2024.09.25.615036

**Authors:** Divyaratan Popli, Benjamin M. Peter

## Abstract

Principal component analysis (PCA) and *F*-statistics are routinely used in population genetic and archaeogenetic studies. Here, we present a statistical framework to combine them into a joint analysis, showing where they coincide, and where slightly different assumptions made can lead to different outcomes. In particular, we discuss the differences of probabilistic PCA, Latent Subspace Estimation and classical PCA, and show that *F*-statistics are more naturally interpreted in a probabilistic PCA framework. We also show that individual-based *F*-statistics can be accurately estimated from probabilistic PCA in the presence of large amounts of missing data. We compare estimates from probabilistic PCA-based framework to ADMIXTOOLS 2 using simulations and published data, and show that this joint estimation framework addresses limitations of estimating F-statistics and PCA independently.

## 1 Introduction

Most populations live in heterogeneous and changing environments, and thus will exhibit some degree of population structure, which we expect to change over time (Charlesworth, 2009; Allendorf, 2017). Over short time scales, there are two principal evolutionary forces at play: First is genetic drift, which increases differentiation between populations over time due to isolation (Song et al., 2006). Second, gene flow between previously isolated populations causes intermediate genotypes and thus reduces differentiation (Lenormand, 2002). For the purpose of this paper, we treat the terms gene flow, admixture and migration synonymously. A common goal in population genetics is to characterize the genetic variation caused by these processes, and to use observed data to infer the processes that caused them (Ellegren and Galtier, 2016).

In particular for humans, there has been a long-standing debate on how we conceptualize genetic population structure, whether populations as discrete units exist, or to what degree they are the result of biased sampling designs (Serre and Pääbo, 2004; Rosenberg et al., 2005; Peter et al., 2020), and how they affect ancestry estimation (Mathieson and Scally, 2020; Simon and Coop, 2023) and impact association studies (Price et al., 2006). Questions like these are of fundamental importance because they impact how we think about race in the context of genetics (Lewontin, 1972; Novembre, 2022), how we measure and provide equitable access to genetic medicine (Popejoy and Fullerton, 2016), and how we conceptualize genetic variation in general.

### 1.1 Overview of methods to study admixture

There is a large array of approaches and methods available to make inferences about population structure (reviewed by Schraiber and Akey, 2015), aiming at different time scales, making different modelling assumptions or treating data differently. One wide class of methods are global ancestry methods. They summarize the entirety of the genome into a small number of summary statistics, with the idea that different genomic loci are (pseudo-) independent replicates of the historical process (Pritchard et al., 2000; Gopalan et al., 2016; Patterson et al., 2012; Alexander et al., 2009; Tang et al., 2005). Global ancestry methods stand in contrast to local ancestry methods, that use an ancestral recombination graph or an approximation thereof, to infer the detailed ancestry of each locus (Lawson et al., 2012; Hellenthal et al., 2014; Speidel et al., 2019; Kelleher et al., 2019). Global ancestry methods are widely used because they tend to be much simpler and easier to interpret than local ancestry methods, and are sufficient for many applications (Pritchard et al., 2000; Patterson et al., 2006).

For large datasets with dozens of individuals spanning a wide range of sampling locations, we can further classify global methods into joint analyses that use data from all individuals in a multivariate framework, such as Principal Component Analysis (PCA) (Cavalli-Sforza and Piazza, 1975; Patterson et al., 2006; Novembre et al., 2008), structure-like methods (Pritchard et al., 2000; Alexander et al., 2009), and summary-statistic-based approaches relying on statistics that include two or a small number of populations (e.g. *F*_*ST*_-based or site-frequency-spectrum based methods), and use large numbers of these summaries to build more complex models (Excoffier and Foll, 2011; Kamm et al., 2020; Gutenkunst et al., 2009).

### 1.2 *F*-statistics

A popular framework based on summary statistics, particularly in the studies of ancient human populations (Orlando et al., 2021), relies on a set of statistics called *F*-statistics (Patterson et al., 2012; Peter, 2016). As we will define in full detail in the theory section, *F*-statistics measure the genetic drift shared between two, three, or four populations (Patterson et al., 2012; Peter, 2016). These patterns of shared drift are then compared to a null model corresponding to a population tree connecting the sampled individuals. Gene flow between distinct populations violates the assumption of treeness, and causes a deviation from the null hypothesis. Thus, *F*-statistics form the basis of an intuitive and powerful framework to test hypotheses of admixture (Fig. 2).

**Figure 1:**
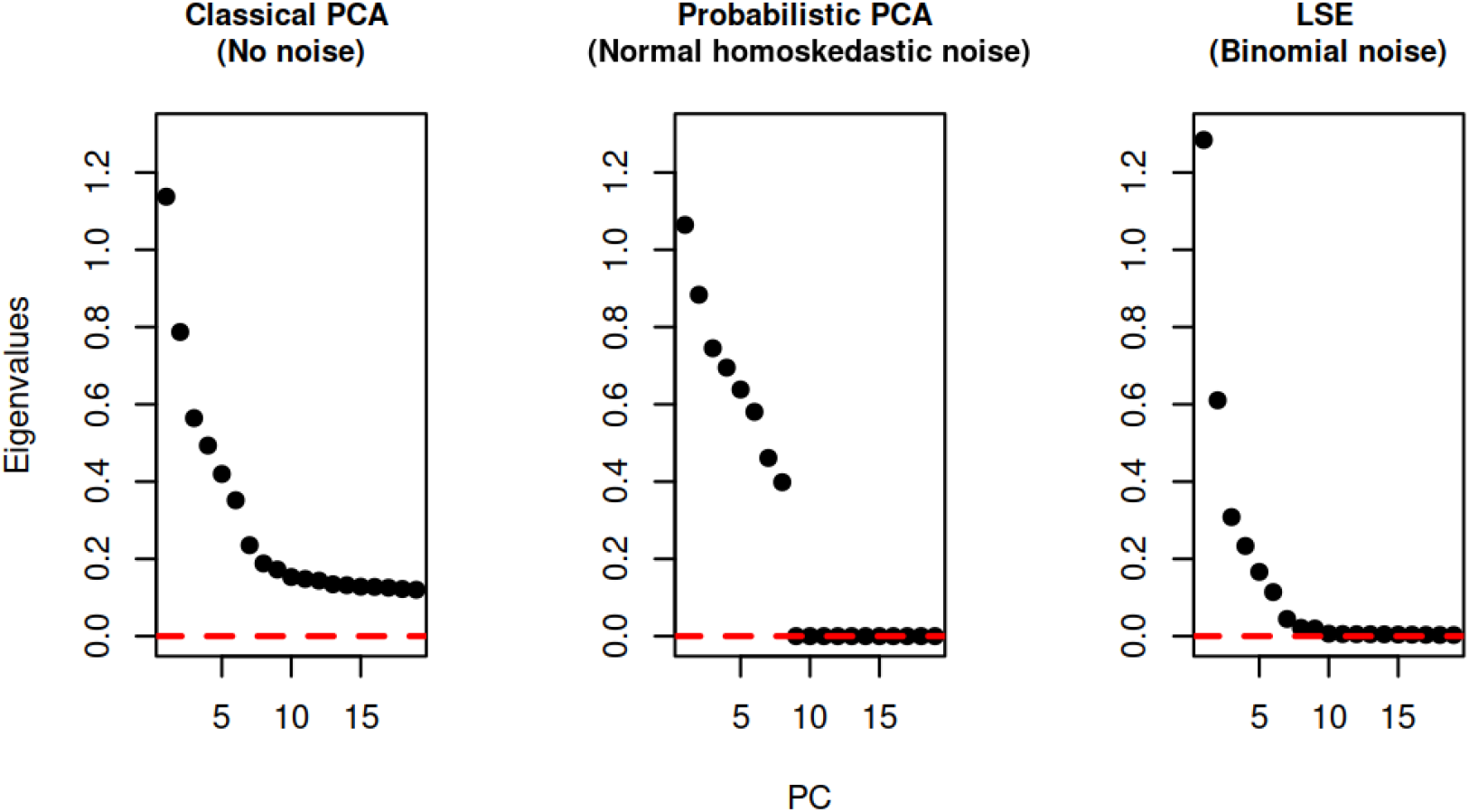
Comparison of classical PCA, PPCA and LSE: we simulated genotypes of 104 individuals belonging to 10 different populations, and compared the top eigenvalues obtained from different PCA methods.

**Figure 2:**
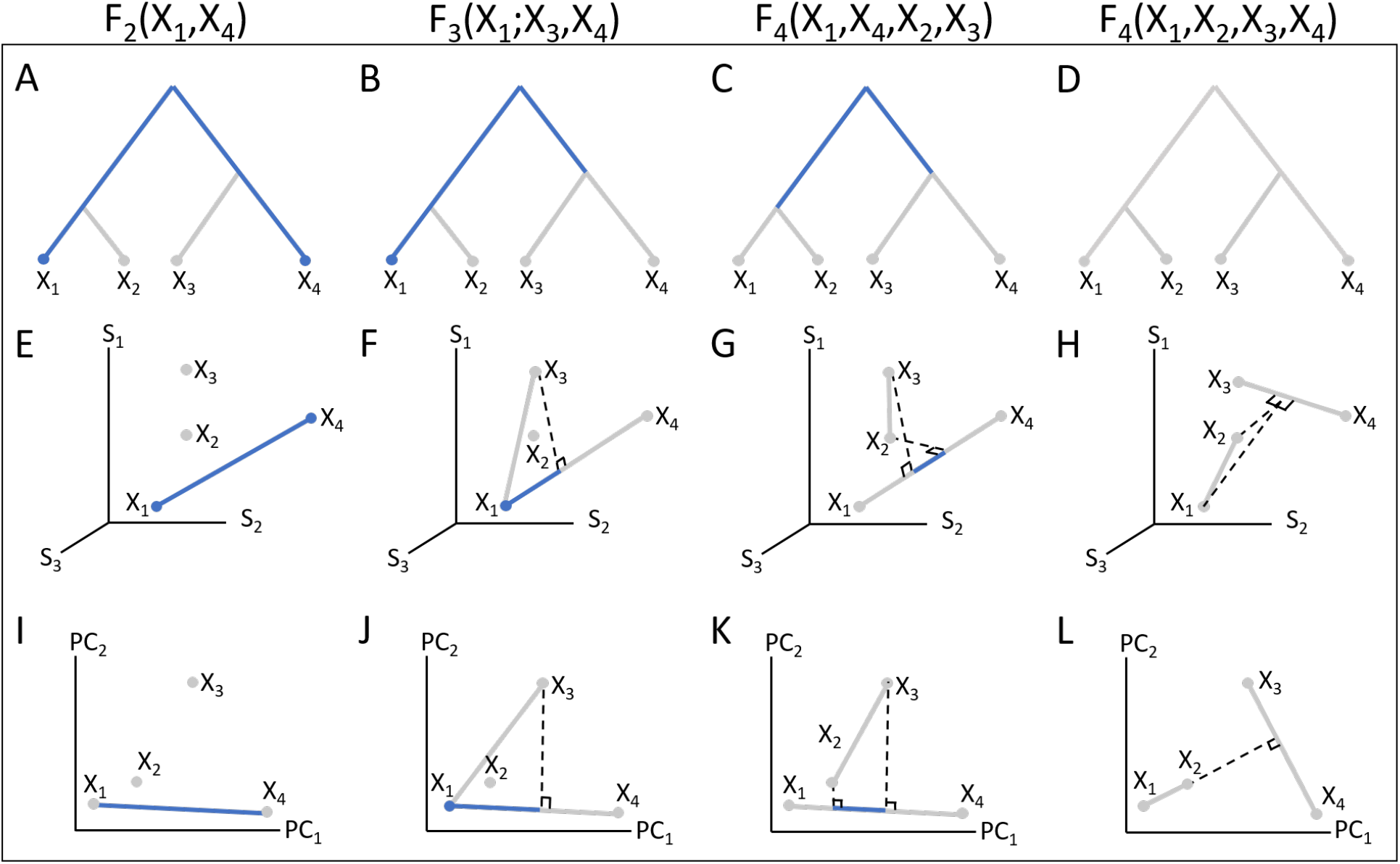
Schematics showing different interpretations of F-statistics. The columns represent *F*_2_(*X*_1_, *X*_4_), F_3_(*X*_1_; *X*_3_, *X*_4_), F 4(*X*_1_, *X*_4_; *X*_2_, *X*_3_), F 4(*X*_1_, *X*_2_, *X*_3_, *X*_4_). The first row shows a tree interpretation of each statistic, the second row shows F-statistics in an allele-frequency space with three axes representing three SNPs, and the last row is the interpretation of F-statistics on a PCA. Blue lines represent the statistic, and the dotted lines represent orthogonal projections. Black squares denote right angles.

Despite their name, *F*-statistics should be thought of as parameters, that are defined in terms of *population* allele frequencies. Since we typically only have genotype data from a small number of individuals, the population allele frequencies are unobserved, and *F*-statistics must be estimated from *sample* allele frequencies. This estimation is not trivial: Patterson et al. showed that a naive estimator would be biased, and introduced a bias-corrected estimator (Patterson et al., 2012). This bias is largest if the sample size is small (e.g. for single genomes), and will reduce in magnitude for larger samples (see eq. 3).

To do inference, we use combinations of statistics that include two, three or four populations. Thus, treating each individual independently would give us the potential to compute more statistics, and thus the highest resolution representation of the underlying population structure. However, large numbers of statistics will be harder to interpret, and would have relatively low statistical accuracy.

Thus, samples are typically grouped into populations a priori, to improve statistical accuracy and to make the interpretation easier. This creates a trade-off between a fine-scale view of population structure with low statistical accuracy, and a coarser-scale representation that has the danger of an overly simplistic view of the genetic structure of the studied species.

This issue is compounded by missing data: In ancient DNA, the amount of preserved DNA is often a limiting factor, and hence low-coverage genomes, and heterogeneous data quality are the norm (Orlando et al., 2021). In studies with dozens of individuals, variable levels of missingness add additional statistical noise that make individual-based statistics even less accurate, leading to a larger pressure to group individuals, and hence simplify the population structure.

To rectify this trade-off between analysis resolution and statistical accuracy, we use the theoretical results linking PCA and *F*-statistics from Peter (2022) to develop a PCA-based framework that jointly estimates *F*-statistics between all sets of individuals in a large data set. This framework has the advantage that it does not require *a priori* assignment of individuals to populations, and allows for the imputation of data missing at random. These advances allow for accurate and finely-grained analyses of population structure.

### 1.3 PCA

PCA is one of the most widely used global ancestry methods to uncover population structure (Cavalli-Sforza and Piazza, 1975; McVean, 2009; Engelhardt and Stephens, 2010). PCA has the advantage that it makes minimal assumptions on the underlying data, and thus can be applied flexibly even when little is known about the underlying patterns of genetic variation. The key empirical feature of PCA is that it tends to cluster similar individuals nearby, and provides an easy to understand and often useful visualization of the genetic variation in the data. PCA is also widely used to model population structure in association studies, where structure is an undesired covariate that has to be regressed out (Price et al., 2006). The big caveat is that it can be difficult to interpret PCAs, since there is no underlying mechanistic model, nor does it lend itself to model comparisons or formal tests of admixture (McVean, 2009; Novembre and Stephens, 2008).

PCA was introduced and popularized as a tool to study human genetic structure by Cavalli-Sforza et al., and using just a handful of genetic loci, they were able to use PCA to accurately describe patterns of human genetic variation, and to make inference about their possible causes, although the way these patterns were analyzed were typically qualitative (Menozzi et al., 1978; Sforza and Sforza, 1995; Cavalli-sforza et al., 1996).

As is still the norm with *F*-statistics, Cavalli-Sforza aggregated individuals into populations before performing PCA. With the availability of genomic data, PCA is now more commonly applied to individual-level data, which results in higher resolution, and does not require grouping of individuals into populations (Patterson et al., 2006; Novembre et al., 2008; Price et al., 2006).

### 1.4 Probabilistic PCA and Latent Subspace Estimation (LSE)

One issue with PCA is that it does not differentiate sources of variation. In population genetic studies, it is often desirable to tease apart the biological variation that is due to the shared population history (leading to the *population* allele frequency), and the statistical variation that is due to missing data, or limited sampling. Analogous to *F*-statistics, applying PCA will result in a biased estimate of population structure when sample allele frequencies are used instead of population allele frequencies. For the remainder of this paper, we will refer to this approach as classical PCA.

Probabilistic PCA (PPCA) is a variant of PCA that aims to rectify this by incorporating an explicit probabilistic framework (Tipping and Bishop, 1999a). The key idea is to decompose the variation in the data into a (low-rank) covariance matrix that models population structure, and a diagonal matrix that captures variation due to sampling.

PPCA was introduced by (Tipping and Bishop, 1999a) in a Gaussian setting, where the sampling error is modelled as normally distributed, and each individual is assumed to have the same sampling error (homoskedastic noise). However, in population genetics, genotypes are discrete and thus the sampling errors are binomially distributed. Furthermore, individuals coming from populations with different effective population sizes will have different heterozygosities resulting in heteroskedastic noise. Thus a more sophisticated model that assumes different binomial sampling noise for each individual has been implemented in Latent Subspace Estimation (LSE) (Chen and Storey, 2015; van Waaij et al., 2023; Cabreros and Storey, 2019). In summary, the main difference between classical PCA, PPCA, and LSE is in the way they model the noise in the observed data due to sampling (see Fig.1). We give a detailed description of classical PCA, PPCA and LSE frameworks in sections 2.2, 2.3 and 2.4 respectively.

### 1.5 Classical PCA and F-statistics

A common analysis paradigm in ancient DNA is to use classical PCA for exploratory and descriptive analyses, and then follow them up with methods based on *F*-statistic for a more formal treatment (typically in a third step, methods that synthesize many *F*-statistics are also applied, although we will not cover those here) (Orlando et al., 2021). Because they were developed independently and occur at different stages of the analyses, classical PCA and *F*-statistics use different data groupings and different normalizations, and are usually not quantitatively compared.

A study by Oteo-García and Oteo (2021) showed that *F*-statistics and the hypothesis testing for admixture can be interpreted geometrically in a multi-dimensional allele frequency space. Since classical PCA only rotates the data points to the axes of maximum variation, Peter (2022) extended this result to show that *F*-statistics can be interpreted geometrically in the context of classical PCA. The study concluded that *F*-statistics and classical PCA are closely related, and *F*-statistics can thus be used to interpret PCA plots; genetic drift will move individuals further apart from each other on a PCA-plot. Independent drift, i.e. unconnected populations, will drift on orthogonal axes in PCA-space. On the other hand, admixed individuals will be placed between their populations of origin, and these placements can be measured using *F*-statistics.

However, while (Peter, 2022) develops a theoretical link between PCA and *F*-statistics, it does not deal with statistical uncertainty, and thus cannot easily be applied to noisy data.

Here, we develop a statistical framework to jointly estimate PCA and *F*-statistics. We show how in particular the choice of the PCA-algorithm (classical PCA, PPCA, LSE) greatly impacts the results. With simulations, we show that *F*-statistics estimated from PPCA-based framework are more accurate compared to that estimated with naive estimators, when there is low sample size and missing data. We also draw comparison between PPCA-based framework and ADMIXTOOLS 2 using published Neanderthal samples. Our approach improves the estimation of *F*-statistics, and leads to some natural suggestions about how and when different PCA methods should be used.

## 2 Theory

In this section, we give a more detailed and formal overview of *F*-statistics and PCA, and their various estimators. We show that the model underlying *F*-statistics is very similar to that of PPCA, and identical to that of LSE. These findings will then be substantiated in the results section using simulated and real data.

### 2.1 *F*-statistics

We follow the original notation of Patterson et al. (2012), and distinguish between the *parameters F*_2_, *F*_3_ and *F*_4_, and their *estimators* from empirical data, denoted by lower-case *f*_2_, *f*_3_ and *f*_4_. The three *F*-statistics are defined in terms of population allele frequencies as follows:

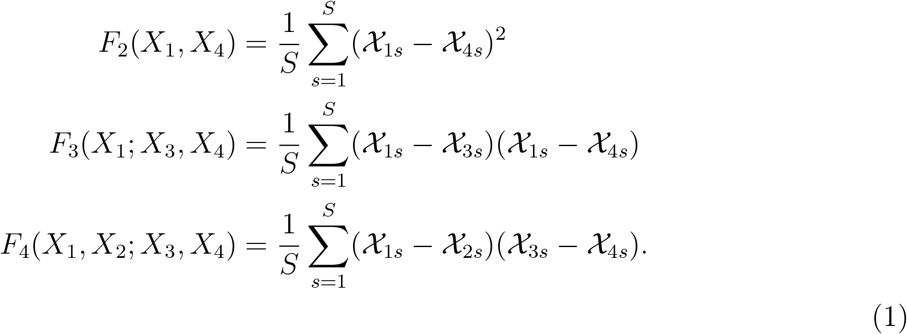

Here, *S* is the total number of SNPs, and *χ*_*is*_ is the (unobserved) population allele frequency in population *X*_*i*_ at SNP *s*.

Assuming a tree-like relationship between populations, *F*_2_(*X*_1_, *X*_4_) is interpreted as the branch length between populations *X*_1_ and *X*_4_ (Fig. 2 A) and it reflects the expected amount of drift that occurred between *X*_1_ and *X*_4_. *F*_3_(*X*_1_; *X*_3_, *X*_4_) represents the amount of drift that occurred on the external branch connecting *X*_1_ to the common ancestor node of *X*_3_ and *X*_4_ (Fig. 2 B). Under a tree-like model, *F*_3_ will always be non-negative. However, in the case where *X*_1_ is admixed between *X*_3_ and *X*_4_, *F*_3_(*X*_1_; *X*_3_, *X*_4_) may be negative, and hence this is used as a test for admixture (Peter, 2016; Patterson et al., 2012). *F*_4_(*X*_1_, *X*_4_; *X*_2_, *X*_3_) represents the covariance between shared drifts between *X*_1_, *X*_4_ and *X*_2_, *X*_3_. This would be represented by the internal branch between the common ancestor nodes of *X*_1_, *X*_2_ and *X*_3_, *X*_4_ (Fig. 2 C). *F*_4_-statistic, with a different permutation of the populations, can be used as test of admixture. *F*_4_(*X*_1_, *X*_2_; *X*_3_, *X*_4_) is expected to be 0 if *X*_1_, *X*_2_, *X*_3_, *X*_4_ are related to each other by a tree (Fig. 2 D). In this case, a significantly non-zero value suggests a departure from the null model of treeness.

It is straightforward to verify that *F*_3_ and *F*_4_ can be written in terms of *F*_2_s as:

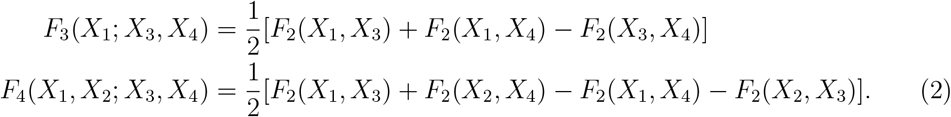

Hence, in practice, all *F*-statistics can be calculated from linear combinations of *F*_2_. Patterson et al. showed that the naive application of eq. 1 to *sample* allele frequency data will be biased, particularly when the sample size is small. They thus introduced the bias-corrected estimator (Patterson et al., 2012):

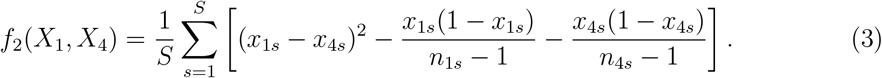

Here, we denote the sample allele frequency for population *X*_*i*_ at SNP s as *x*_*is*_, and the number of non-missing haploids in population *X*_*i*_ at SNP s as *n*_*is*_. When only a single (diploid) individual is sampled from population *X*_*i*_, *n*_*is*_ = 2 and so this equation still works in this case. However, for (pseudo-) haploid samples *n*_*is*_ = 1 and the denominators are zero. Thus, the unbiased estimators do not exist for single pseudohaploid samples.

Using eq. 2, we can see that this sampling bias also affects the calculation of *f*_3_, but the correction terms cancel out for *f*_4_, and the equations for *f*_4_ and *F*_4_ coincide:

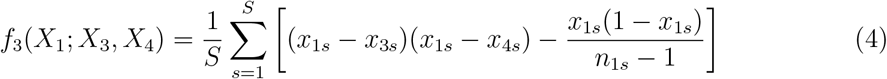

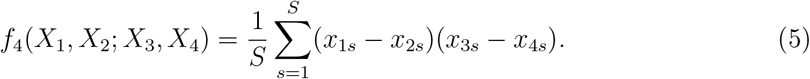

Oteo-García and Oteo (2021) showed that *F*-statistics can be defined more generally in a geometrical framework. In this framework, each population can be represented as a point in a high-dimensional allele frequency space. *F*_2_(*X*_1_, *X*_4_) is then the squared Euclidean distance between points representing populations *X*_1_ and *X*_4_ (Fig. 2 E). *F*_3_(*X*_1_; *X*_3_, *X*_4_) is then a dot product of the vectors 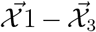 and 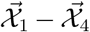 (Fig. 2 F). *F*_4_(*X*_1_, *X*_4_; *X*_2_, *X*_3_) is a dot product of vectors 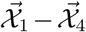 and 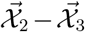 (Fig. 2 G), and *F*_4_(*X*_1_, *X*_2_; *X*_3_, *X*_4_) is a dot product of vectors 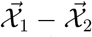 and 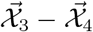 (Fig. 2 H). We define the vector of allele frequencies of population *i* as 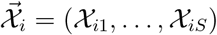.

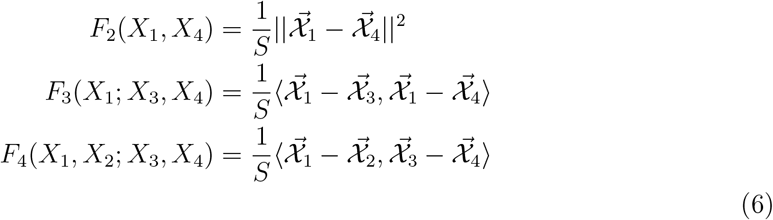

Crucially however, these interpretations only hold for the (unobserved) population allele frequencies, but not for the (observed) sample allele frequencies and thus, they cannot be directly applied to noisy data.

### 2.2 Classical PCA and F-statistics

This geometric framework provides a complementary way to understand the properties of *F*-statistics (Oteo-García and Oteo, 2021). However, many population genetic studies use a large number of SNPs (in the order of a million), and it is not possible to visualize population vectors in such a high dimensional space. Peter (2022) showed that one can do dimensionality reduction on such datasets with classical PCA, and use the top PCs to estimate *F*-statistics efficiently.

We illustrate this in Fig. 2: *F*_2_(*X*_1_, *X*_4_) can be thought of as the squared Eucledian distance between populations *X*_1_ and *X*_4_ in PCA-space. Similarly, *F*_3_(*X*_1_; *X*_3_, *X*_4_) is represented as the length of projection of the vector *X*_1_ *− X*_3_ on *X*_1_ *− X*_4_. Internal branch length *F*_4_(*X*_1_, *X*_4_; *X*_2_, *X*_3_) can be described as the length of projection of *X*_2_ *− X*_3_ on *X*_1_ *− X*_4_ on PCA, and the test of admixture *F*_4_(*X*_1_, *X*_2_, *X*_3_, *X*_4_) is equivalent to the length of projection of *X*_1_ *− X*_2_ on *X*_3_ *− X*_4_. See Peter (2022) for full details.

To formalize the relationship between *F*-statistics and PCA, let us assume our dataset **X** has M individuals and *S* SNPs, such that our allele frequency matrix **X** has the dimension [*M* × *S*]. The *i*-th row of **X** corresponds to the *S*-dimensional row-vector 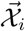 whose entries are allele frequencies ∈ [0,1]. Classical PCA of mean-centered **X** allows us to project this *S*-dimensional data onto a *q* dimensional subspace, where *q* < *M. q* = *M −* 1 represents the case where we retain all the PCs, and thus classical PCA only rotates **X**. However, in practice we often only need few PCs (*q ≪ M*) to explain most of the variation in the genetic data (Peter, 2022), which can greatly simplify calculations and visualizations.

A common algorithm to estimate PCs is via Singular Value Decomposition (SVD). For this approach, we first mean-center **X** to a matrix **X**_*c*_ by subtracting the mean genotype *μ*_*s*_ from all the rows of **X**, and then decompose **X**_*c*_ into an orthonormal matrix **U**, a diagonal matrix Λ, and another orthonormal matrix **V**^**T**^.

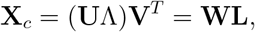

We can perform SVD to decompose **X**_*c*_ into a product of **W** = **U**Λ and **L** = **V**^*T*^. In the context of PCA, **W**_[*M×M*]_ is a matrix of principal components (where the *i*-th column corresponds to the *i*-th PC) and contains information about structure, while **L**_[*M×S*]_, also known as the SNP loadings contains the contribution of each SNP to each PC (Gower, 1966), and can be used to identify outlier SNPs that may be potential candidates for selection (Meisner et al., 2021a).

Since *F*-statistics can be written as dot products in an allele frequency space (eq. 6), and dot products are invariant to rotation, classical PCA will not change *F*-statistics as long as we retain all PCs and we can calculate *F*-statistics from the PCs directly (Peter, 2022):

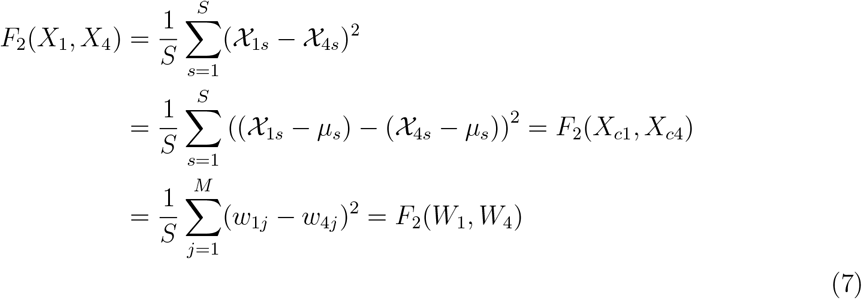

A difficulty in the practical application of this result is that the geometric considerations of Oteo-García and Oteo (2021) and Peter (2022) only hold for the (generally unobserved) population allele frequencies, but not for sample allele frequencies. In ancient DNA, PCA is most commonly run directly on individual-level genotype data (Patterson et al., 2006), and hence on the biased sample allele frequencies. Thus, applying the classical PCA-based estimator (eq. 7) to calculate *F*-statistics would likewise result in biased estimate.

Oteo-García and Oteo (2021) resolved this by calculating *F*-statistics using populations with large number of individuals and with no missing data. In Peter (2022), unbiased estimates of the PCA reconstructions were obtained indirectly by first calculating all pairwise *F*_2_-statistics, and then performing a multidimensional-scaling decomposition equivalent to population-level PCA.

### 2.3 PPCA and F-statistics

Since PPCA and LSE separate the population variation from sampling uncertainty, we can use them to calculate (approximately) unbiased *f*-statistics.

Here, we implement approaches based on PPCA and LSE that aim to calculate the bias-corrected estimates of *F*_2_ from PCA, by explicitly separating out the sampling error.

The first approach is based on probabilistic PCA (PPCA) (Tipping and Bishop, 1999a; Agrawal et al., 2020). The simple idea here is to get the covariance matrix for all pairs of samples, along with an error term. We do this by estimating matrix **W**, such that **WW**^*T*^ denotes the covariance matrix, and Ψ represents the average covariance due to sampling noise. Then, the model fit by PPCA can be written as:

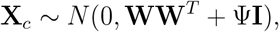

where *N* denotes a multivariate normal distribution, **X**_*c*_ is again the centered genotype matrix, **W** is a *M × q* matrix of linear mappings, **I** is the identity matrix and Ψ is a scalar noise term. Intuitively, **WW**^*T*^ captures the covariance in the observed data, analogous to the *F*-statistics, and Ψ**I** is analogous to the bias-correction term. Since Ψ is a scalar, all entries on the diagonal of Ψ**I** will be the same, and thus the model is homoskedastic, i.e. all individuals are assumed to have the same error.

To fit the PPCA model to a dataset, we need to estimate the parameters of the model, namely **W** and Ψ, given the observed data. For complete data, Ψ and **W** can be calculated from SVD using the maximum-likelihood estimators (Tipping and Bishop, 1999b):

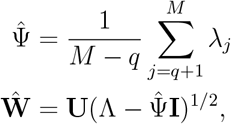

where λ_*j*_, the *j*-th eigen value, is the *j*-th entry on the diagonal of Λ.

Intuitively, the error term is just an average of all the eigenvalues that correspond to the eigenvectors that are thought of as contributing to the sampling noise, and therefore it is subtracted from the diagonal matrix of eigenvalues Λ. Thus, the MLE solution of PPCA results differs from that of classical PCA only in that the first *q* PCs are “shrunk” using a common term that incorporates the noise discarded in the PCA, and the last *M − q* PCs are set to zero. Notably a classical PCA plot would look almost the same, only the axes scales would be different. Notably, for PPCA we need to set the number of retained dimensions *q* a priori, since changing *q* requires rescaling all PCs. In addition, setting *q* = *M −* 1 or Ψ = 0 recoups classical PCA. Similar to the case of classical PCA, we can use eq. 7 to calculate *F*-statistics using PPCA.

### 2.4 LSE and F-statistics

The drawback of PPCA is that it models the sampling error as the same for each sample, but in genetic data the error is binomially distributed. In addition, one needs to know *q* a priori. LSE (Latent Subspace Estimation) is a dimensionality reduction technique that addresses these issues. LSE is quite similar to PPCA, with the difference that it accounts for the heteroscedasticity in the data (Chen and Storey, 2015), and explicitly models the binomial error in genetic data. The intuitive idea here is to calculate the sampling error for each sample assuming a binomial distribution, and to remove this error from the covariance matrix before doing SVD. So, in this algorithm, we calculate the heterozygosity 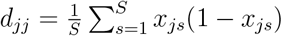 from **X**. We define **D** as a diagonal matrix with entries *d*_*jj*_. One can then estimate covariance matrix 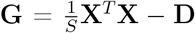. The eigenvectors of **G** then span the latent subspace of **L**, and the expected value of the smallest *M − q* eigenvalues converge to 0 for large M (Cabreros and Storey, 2019).

Crucially, if we use *all* the PCs, the *f*-statistics coincide with LSE (Fig. S4, see the appendix (section 6) for a derivation):

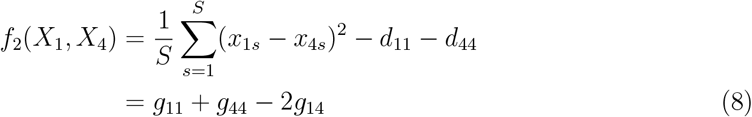

The covariance used by LSE is an estimated covariance matrix (van Waaij et al., 2023), and its properties differ from the sample covariance matrix used in classical PCA. In particular, sample covariance matrices are positive semi-definite, which means that all eigenvalues are non-negative. This ensures that the eigenvalues represent the variance among the individuals along the corresponding PCs. In contrast, for an estimated covariance matrix, the expectation of the smallest *M − q* eigenvalues is zero, and thus an unbiased estimate will have both positive and negative eigenvalues. Therefore, in case of LSE, we can calculate *F*_2_ by adjusting eq. 7 to include the sign of the eigenvalues:

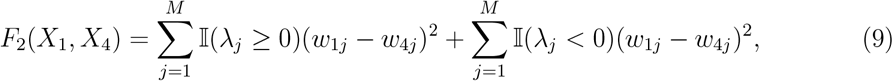

where λ_*j*_ denotes the *j*-th eigenvalue of **G**, and 𝕀 denotes the indicator function that is 1 when the condition is satisfied, and -1 otherwise.

### 2.5 Missing data

It can be difficult to estimate *F*-statistics when there is high amount of missing data. A conservative approach to estimate individual-based *F*-statistics is to only retain sites where data is present from every single individual in the data set. However, even for moderately large data sets that quickly becomes prohibitive: As a toy example, consider a data set with 100 (haploid) individuals with 10% missing data, and 1,000,000 sites. Out of those, only 26 are expected to be covered in every single individual, which makes this approach not feasible.

Thus, grouping individuals into populations can dramatically increase the number of sites retained. A common practice is to retain sites if at least one individual in each population has data. In the above example, if we grouped the 100 individuals into 10 populations of 10 individuals each, and retained sites where at least one individual in each population carried data, we would expect no missing sites in almost all cases. However, grouping individuals may not be justified when they do not form discrete clusters, or when there are very few samples whose population assignments are known.

Missing data is also a challenging problem while computing PCA, since SVD and eigendecompositions cannot be performed with missing data. One way to implement a PCA in case of missing data is to start with assigning the mean genotype to the missing values (mean imputation), and then perform PCA to reconstruct or impute the missing values, and keep iterating until convergence. This can be done both for classical PCA (Meisner et al., 2021b) and for PPCA (Tipping and Bishop, 1999a).

Another popular way to deal with missing data is to first construct a PCA using samples that have no missing data, and then project the low-coverage samples on this PCA (Patterson et al., 2006; Price et al., 2006). Projection is a useful way to check if the low-coverage samples from the same population fall into the same position as the high coverage ones. In this approach, one needs to make sure that the construction of PCA is not affected by sampling bias (such as in the case of classical PCA). However, since sampling noise is independent for each projected sample, projection of samples on a constructed PCA does not require additional modeling of sampling bias.

## 3 Results

In this section, we first describe the coalescent simulations we use to study the theoretical and statistical properties of different PCA methods. We compare the *F*-statistics predicted from different versions of PCA to those calculated using ADMIXTOOLS 2 (Maier et al., 2022). Finally, we illustrate a practical application of our approach on a dataset of Neandertal genetic variation. We developed a snakemake pipeline (Mölder et al., 2021) with python and R scripts for all the analyses described here (see Github (Popli, 2024) or section 4 for details).

### 3.1 Evaluation on simulations

We simulated 4 populations (*X*_1_, *X*_2_, *X*_3_, *X*_4_) with 11 individuals each and 6 populations (*X*_5_,…, *X*_10_) with 10 individuals each using slendr (Petr et al., 2022). In these simulations, populations *X*_6_ to*X*_10_ were created from pairwise admixture events between populations *X*_1_,..,*X*_5_. In addition, we simulated a unidirectional gene flow from *X*_3_ to *X*_2_ with varying migration rates (0%, 1%, 5%) (see Fig. 3). We generated 20 simulations for each case of migration rate to evaluate our framework. In these simulations, we used a mutation rate of 10^*−*8^ per base per generation, a recombination rate of 10^*−*8^ per base per generation, and a generation time of 30 years. The effective population size for each population was set to 1000.

**Figure 3:**
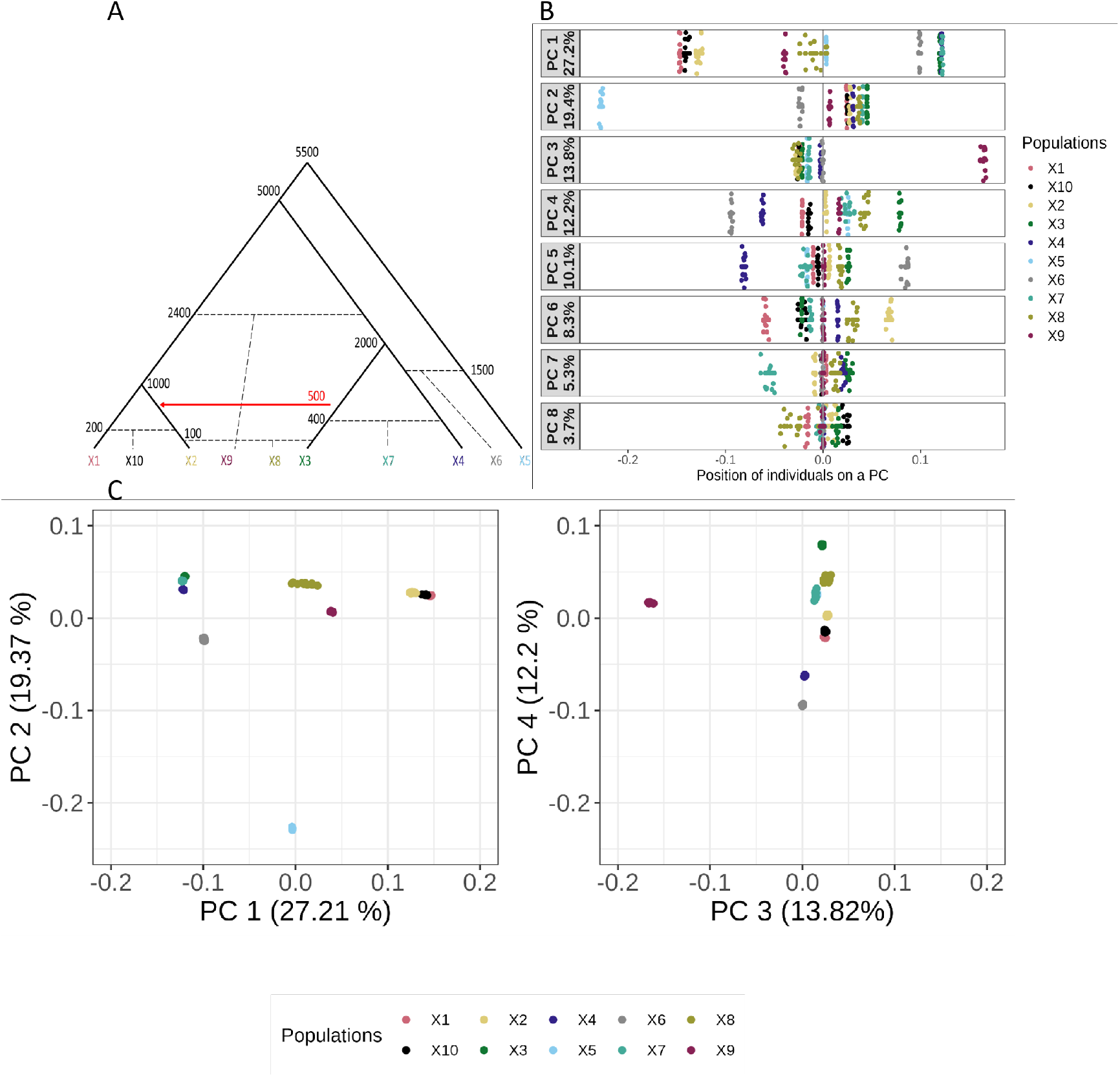
Overview of simulations. **A.** Schematic of the simulated model. Numbers reflect split, migration and admixture times (in generations). Admixture events are shown with dashed lines, and the red arrow represents a unidirectional migration from *X*_3_ to *X*_2_. **B**. PPCA (with 8 PCs) of data simulated using the admixture graph depicted in A. Percentage labels show variation explained per PC. **C**. PPCA-biplots of the first 4 PCs.

### Interpretation of PCA of admixture graph

The admixture graph in Fig. 3 A shows a schematic of the population structure and demographic history for simulations. We can visualize this structure using two different PPCA plots: first we plot the first eight PCs on the same scale (Fig. 3B), and we also show the more standard PPCA biplots (Fig. 3C). The genetic structure generated by the admixture graph is apparent in both visualizations: For example, the first PC mainly separates the “left” clade (with *X*_1_,*X*_2_ and *X*_10_), from the “right” clade (*X*_3_,*X*_4_ and *X*_7_). The outgroup (*X*_5_), is placed very close to zero here, which is because PC1 mainly reflects the variation between these clades, and the outgroup branch leading to *X*_5_ is orthogonal to that variation.

PC2, on the other hand, primarily separates out the outgroup (*X*_5_) from the remaining samples, with the admixed population *X*_6_ falling in between.

Thus, even though *X*_5_ is the most genetically diverged population, this is not evident from PC1. The reason for this is that PCA maximizes the sum of all pairwise *F*_2_-distances between individuals, and hence the ordering of PCs depends on sample configuration (McVean, 2009; Elhaik, 2022). In our case, there are only ten individuals in *X*_5_, whereas we have a total of 84 individuals in the other populations (all except *X*_6_, which has ancestry from *X*_5_). In this case, the sum of the 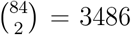 pairwise *F*_2_-distances within the clade is larger than the sum of the 84 *×* 10 = 840 distances between individuals from the clade to *X*_5_.

Next, PC 3 shows the axis of variation between *X*_9_ and the other populations. Due to the drift on *X*_9_ for many generations, it is hard to detect the signal for admixture (*f*_3_(*X*_9_; *X*_1_, *X*_4_) = 0.0008). In comparison, the admixture between *X*_1_ and *X*_2_ leading to *X*_10_ is detectable (*f*_3_(*X*_10_; *X*_1_, *X*_2_) = −0.0002). *f*_3_ is calculated with the top 8 PCs in the above cases using eq. 7. The two next PCs, PC4 and PC5 mainly model the variation with the “right” clade, and PC6 splits up *X*_1_ and *X*_2_ within the “left clade”. In all three of the PCs, populations outside the clades plot very close to zero, which is because in a tree, the within-clade variation is independent of that of the clade to individuals outside of it (Felsenstein, 1973).

Thus, we find that for tree and admixture graph models, most PCs tend to show variation within a single clade, and we need a fairly large number of PCs to tease out the full tree structure. This is consistent with the theoretical expectation that the covariance matrix for a (population) tree has full rank (Felsenstein, 1973), i.e. we would expect the number of PCs required to reflect tree structure to be on the same scale as the number of populations (Patterson et al., 2006).

PCA is most commonly visualized in biplots (Fig 3C). A main advantage is that this provides a 2D-visualization of the high-level population structure (in our case, the separation of the outgroup from the ingroup samples and between the two main clades). However, we lose the structure within each clade, that is represented by PCs 3 and higher. Plotting a second biplot of PC3 vs PC4 is somewhat less informative, because the variation already explained by the higher PCs is absent.

In contrast, plotting PCs separately on one dimension has the advantage that the orthogonality of the PCs becomes more apparent, and also allows for easier quantitative comparison on how much each PC explains: The spread (i.e. variance explained) by each PC gets narrow and narrower as we move to higher PCs, and we can easily plot a large number of PCs. The drawback of this representation is that the correlation and 2D-structure in PC-space gets de-emphasized. We plot the first 10 PCs from PPCA (scale=8), classical PCA and LSE (Fig. S1, S2, S3), and find that the population structure represented by the PCs does not change. However, the PCs are scaled differently for different method, and so the absolute position of individuals varies. In particular, for PPCA (scale=8), all PCs > 8 are scaled with zero, and hence PC 9 and PC 10 in this case look very different from that of classical PCA and LSE. It is interesting to note that a single PC is not informative. One needs to look at all the PCs to get a sense of the population structure. However, the information in the tree and the PC plot is the same, and so the tree can be regenerated from the (top) PCs.

#### PCA-based *f*_2_ estimates

We use eq. 7 to write python scripts for calculating *f*-statistics via classical PCA, PPCA and LSE, and to compare these *f*-statistics to the true values obtained from the simulations. We implemented PPCA and LSE using R scripts, and classical PCA using prcomp function in R (Popli, 2024).

To compare how the different *f*_2_-estimates behave, we calculate *f*_2_(*X*_1_, *X*_4_) with an increasing number of PCs for the three methods (Fig. 4).

**Figure 4:**
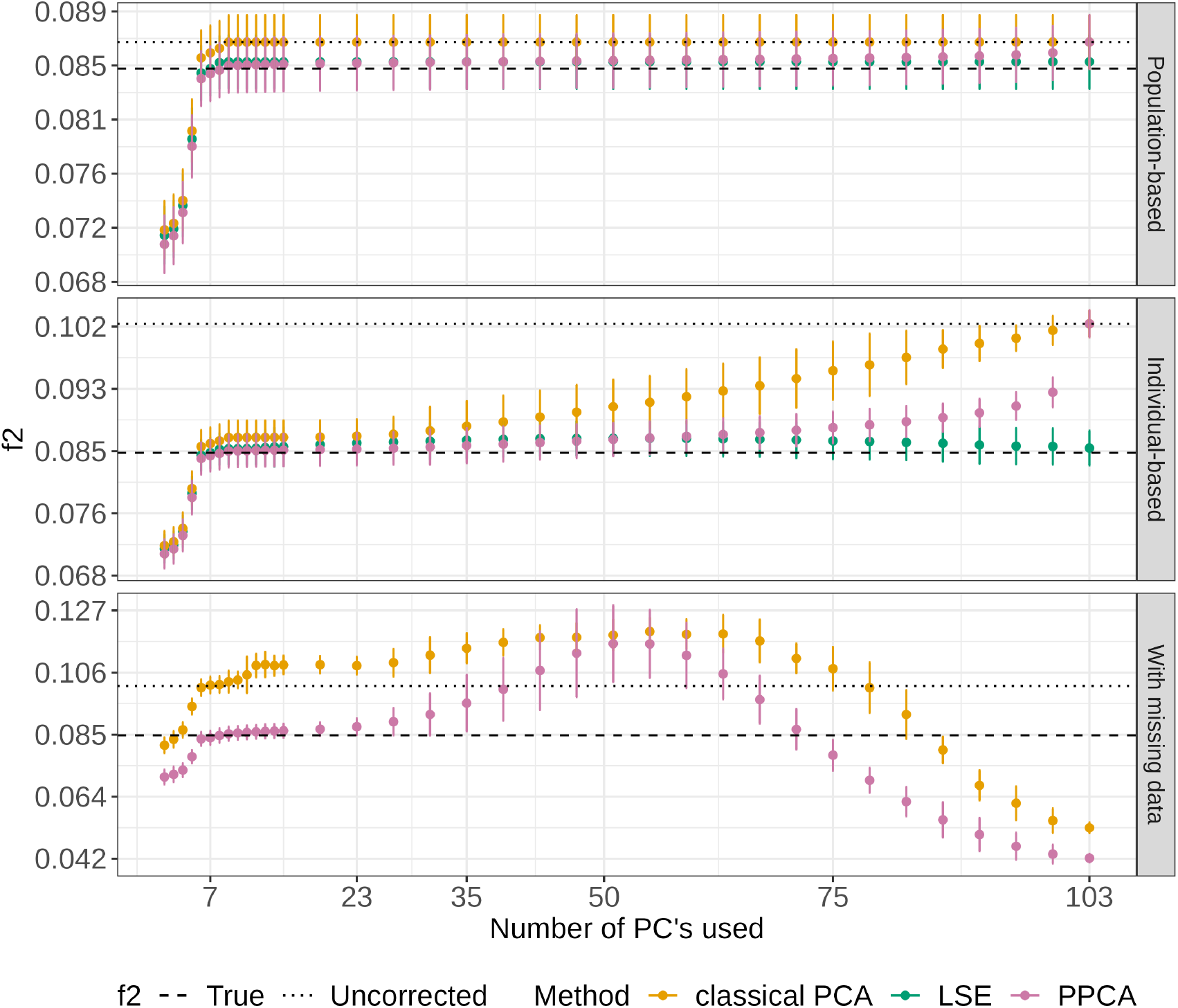
Comparison of PCA approaches using *f*_2_(*X*_1_, *X*_4_) estimated using 11 individuals for each population (top), 1 individual for each population (middle), and 1 individual for each population with 50% missing genotypes (bottom). Dotted line represents biased estimate and dashed lines show the true value of *F*_2_ from the simulations. Error bars represent 2 standard errors. The bottom panel includes only PCA and PPCA, since LSE implementation with missing data is outside the scope of this paper.

In the top panel of Fig. 4, we use eleven diploid individuals per population. Since the sampling error is low with eleven individuals, the bias-correction is small, and the biased *f*_2_-estimate (0.0866, calculated using eq. 1) is similar to the true value calculated from branch lengths (0.0845). In contrast, the bias is much larger when using a single diploid individual (middle/lower panel), with the biased estimate at 0.102.

In either case, we find that classical PCA and PPCA both converge to the biased estimate with increasing number of PCs, whereas LSE converges to the unbiased statistic, as expected from eq. 8. However, the rates of convergence differ: For population-based samples, classical PCA needs just ≈ 7 PCs to be very close to the biased estimate (0.0861), and the last 94 PCs only provide a marginal change. In contrast, in the individual-based analysis the estimate reaches a plateau between 7 ∼ 23PCs where its value is just a slight overestimate of the true value (0.0861 at 7 *∼* PCs, 0.0868 at 23 PCs).

For PPCA, the number of PCs is the rank of the covariance matrix estimated, with the rest of the variation explained by the noise term. In the individual-level analysis, the lower impact of the number of PCs for PPCA becomes apparent *f*_2_ = 0.0841, 0.0850, 0.0863 using 7 PCs, 23 PCs and 51 PCs respectively, demonstrating that the PPCA-derived *f*-statistics are very robust to the number of PCs used. The third method, LSE, converges to the true value when all PCs are used. However, similar to PPCA we find that using 7, 23 or 51 PCs yields estimates with a minimal bias (*f*_2_ = 0.085, 0.0859, 0.0865, respectively). It is interesting to note that the *f*_2_ estimated with LSE initially increases with additional PCs, and then reduces to converge to the true value. This is because many small eigenvalues are negative and therefore *f*_2_s calculated from the corresponding PCs are subtracted (see section 5.0.2).

In the final panel, we estimate *f*_2_ with data where we introduce 50% missingness (Fig. 4). Implementing LSE with missing data is beyond the scope of this paper, and we only compare PCA and PPCA. We again find that PPCA performs well over a fairly large range of number of PCs used but that the estimates become more erratic beyond 23 PCs (*f*_2_ = 0.0836, 0.0874, 0.1158 using 7, 23 and 51 PCs respectively). In contrast, the estimates from classical PCA stay close to the biased *f*_2_-statistic between 7 to 23 PCs.

### 3.2 Estimating F-statistics using PCA

We next compare PCA-based estimates of *F*-statistics to those computed using ADMIXTOOLS 2 (Maier et al., 2022), which is a state-of-the-art re-implementation of ADMIXTOOLS (Patterson et al., 2012) that gives equivalent results. As we have shown, estimates of *F*-statistics based on classical PCA are biased, and so we omit it from this analysis. We compare PPCA and LSE with 8 and 12 PCs, using two numbers because the number of PCs to use will not be known in most applications. We first compare *f*_2_, *f*_3_ and *f*_4_ estimated by these methods from complete simulation data, with all individuals in each population, and no missing genotypes. In this case we find that all the methods perform well, and result in *F*-statistics that are very close to the truth (Fig. S5).

We then subsample one individual from each population, and observe that in this case both PPCA and LSE-based frameworks perform at least as good as ADMIXTOOLS 2 (Table S1, Fig. S6) and the mean estimate from each method is close to the true value. However, the error bars for *f*_2_ estimates (obtained using the point estimates from 20 simulations) are slightly lower for PPCA (scale=8) compared to those of ADMIXTOOLS 2: For *F*_2_(*X*_1_, *X*_2_), *F*_2_(*X*_3_, *X*_4_), *F*_2_(*X*_2_, *X*_3_) and *F*_2_(*X*_1_, *X*_4_), ADMIXTOOLS 2 standard errors are 0.0018, 0.0015, 0.0027, and 0.0012 while the PPCA and LSE-based estimates (scale=8) for the same statistics are 0.0012,0.0012,0.0024,0.0011 and 0.0012,0.0012,0.0025,0.0011 respectively). The improved accuracy of PCA-based tools versus ADMIXTOOLS 2 is explained because PCA incorporates a succinct summary of the full data of all the individuals, and thus the PCA-based estimates can “borrow” information from related individuals in the sample that are not used to calculate the statistic at hand. In contrast, ADMIXTOOLS 2 has only one individual from each population to assess structure / admixture, and while the estimates based on ADMIXTOOLS 2 are minimum-variance estimators for these subsets of the data (Patterson et al., 2012), PCA-based methods do better whenever we have data from additional individuals.

### 3.3 Missing data

Next, we address the issue of missing data and evaluate the estimation of *F*-statistics using one individual from each population with these methods when there is 50% random missing data. Our implementation of PPCA with missing data is inspired from EMU (Meisner et al., 2021b), and is described in Methods section 4.1. However, the EMU implementation aims to reproduce classical PCA, and hence would result in biased estimates of *F*-statistics. Our correction is able to produce accurate *f*-statistics using PPCA (Fig. 5, Table S1, whereas both EMU and ADMIXTOOLS 2 estimates are inflated .For the case of missing data, ADMIXTOOLS 2 was run with maxmiss=1, so no SNPs are excluded. It is not possible to run ADMIXTOOLS 2 using sites with no missing data since with 50% missing sites, very few sites are without missingness.

**Figure 5:**
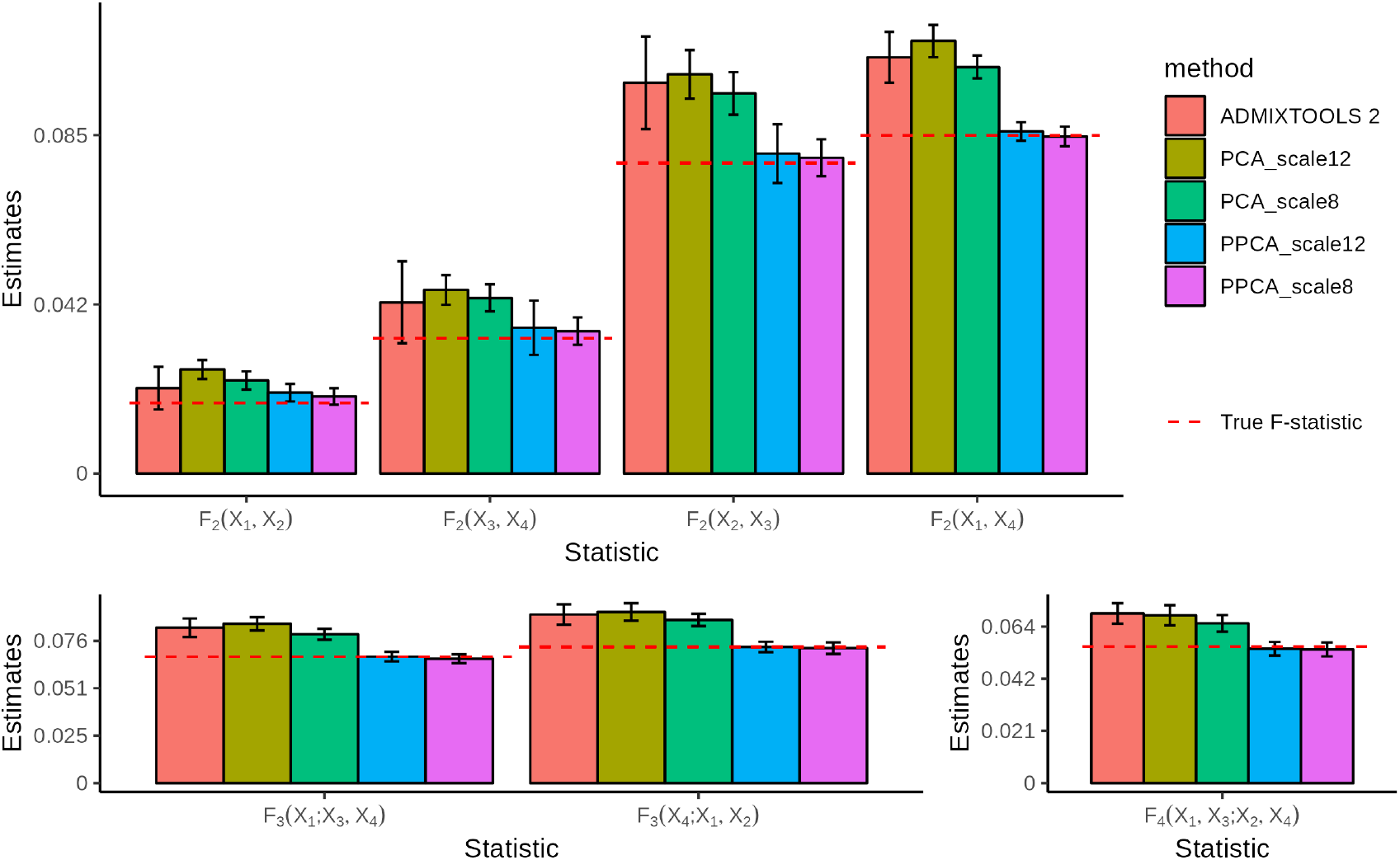
Comparison of PPCA and classical PCA to ADMIXTOOLS 2 in the presence of 50% random missingness, using genotypes from one individual from each population. Red dashed line reflects the true value. Error bars represent 2 standard errors.

**Figure 6:**
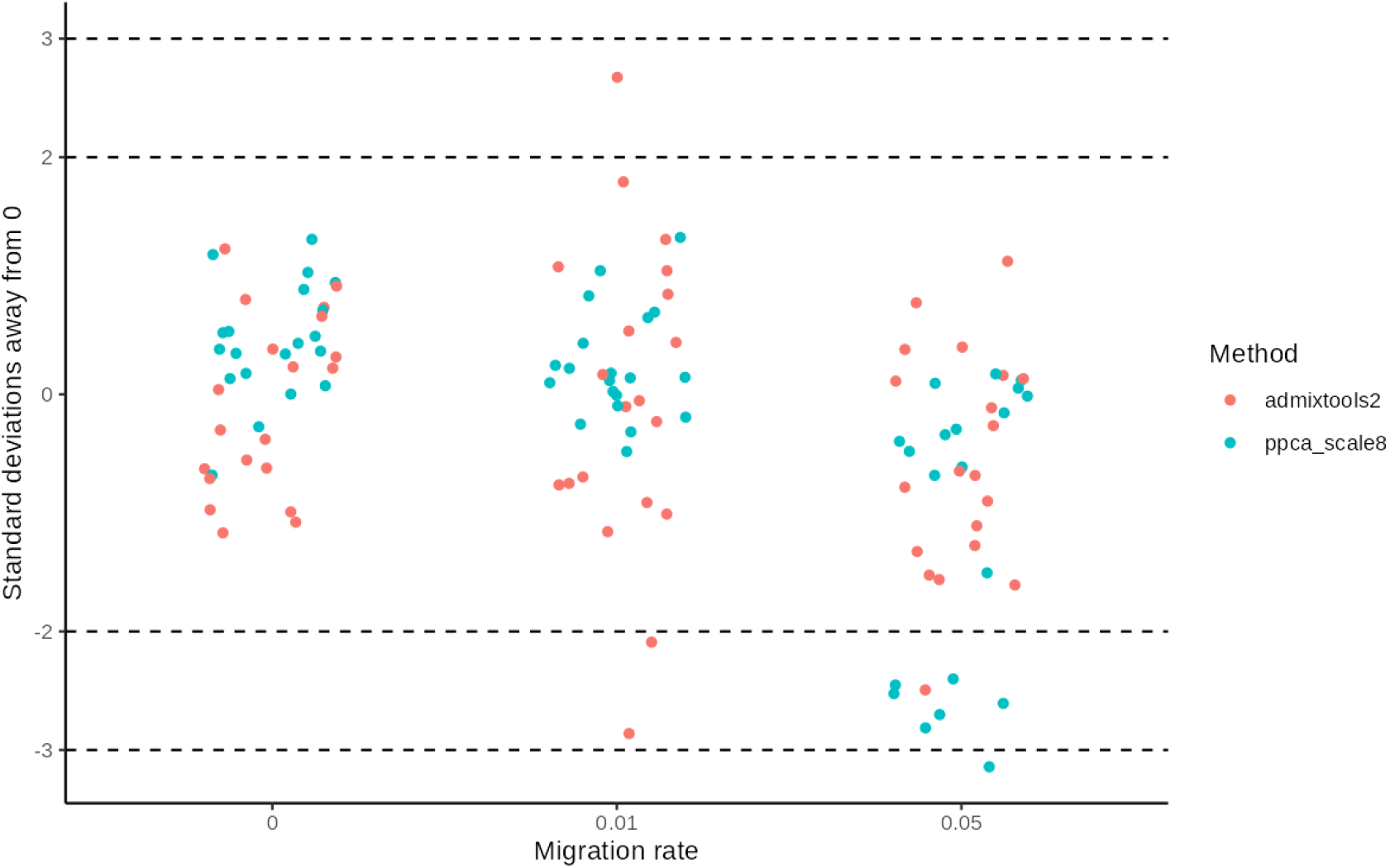
Test for admixture with individual-based *f*_4_(*X*_1_, *X*_2_, *X*_3_, *X*_4_) statistic. We compare ADMIXTOOLS 2 (orange) to PPCA-based-framework (blue) using 20 simulations with 50% missing genotypes. Jitter is added on x-axis for visual clarity.

### 3.4 Test of admixture

A major application of *F*-statistics are tests of admixture (Orlando et al., 2021). We showed in the previous section that PPCA framework can be used to calculate the point estimates of *F*-statistics. In this section we show that we can also get standard errors for these estimates using block-jackknife (Kunsch, 1989; Maier et al., 2022; Patterson, 2020), and use these to do hypothesis testing for admixture. We simulate a gene flow from *X*_3_ to *X*_2_ 500 generations ago with the migration rate of μϵ[0, 0.01, 0.05]. We then compare the estimates of *F*-statistics from PPCA framework to that from ADMIXTOOLS 2. We first test for admixture by checking if the estimate of *F*_4_(*X*_1_, *X*_2_; *X*_3_, *X*_4_) is significantly different from 0. We show that when there are 11 individuals in each population, both methods perform well (Fig. S7). In case of 0 migration rate, both methods estimate *F*_4_ for all simulations to be close to 0, while at 5% migration rate, ADMIXTOOLS 2 and PPCA-framework have the power to detect admixture (with *F*_4_ estimate 2 standard deviations below 0) in 90% and 70% simulations respectively. At migration rate of 1%, both the methods are unable to find admixture between *X*_2_ and *X*_3_, and instead incorrectly predict admixture between *X*_1_ and *X*_3_ (*F*_4_ estimate is 2 standard deviations above the mean) for one simulation. Reducing the number of individuals to 1 from each population and adding 50% missingness reduces the power for both the methods. In this case, both methods have no false positives in case of 0 migration rate, and at 5% migration rate ADMIXTOOLS 2 and PPCA framework detect admixture in 5% and 35% simulations respectively. At 1% migration rate, ADMIXTOOLS 2 infers admixture between *X*_2_ and *X*_3_ in 2 simulations and incorrect admixture between *X*_1_ and *X*_3_ in one simulation, while PPCA framework shows no prediction of admixture.

### 3.5 Evaluation on a Neandertal dataset

To test our framework on real data, we use it to re-analyze a published dataset of archaic humans from Eurasia (Hajdinjak et al., 2018). This dataset consists of low-coverage sequence data from late Neandertals from Western Eurasia, which were included as pseudo-haploid genotypes. For context, the authors also added the high-coverage diploid genotype sequences from Vindija cave, Croatia (Vindija33.19)(Prüfer et al., 2017) and Densiova Cave, Russia(Altai, Denisova)(Prüfer et al., 2014; Meyer et al., 2012). The Denisova specimen (*Denisova 3*) was the first Denisovan discovered, whereas the Altai Neandertal (*Denisova 5*) is a Neandertal from the same site that has been shown to be on a different branch than all the late Neandertals (Prüfer et al., 2014, 2017). We find that the results of PPCA and classical PCA with projection, as implemented in smartPCA (with only two PCs to make our analysis same as the original study), provide a visually very similar PCA plot (Fig. 7). In both cases, PC1 separates out the Denisovan from the Neandertals, whereas PC2 results in a gradient from Altai to *Vindija 33*.*19*. The main difference is that because classical PCA includes sampling noise in the analysis, it overestimates the absolute magnitude of differentiation; classical PCA visualizes the biased estimator for *F*_2_, wheras PPCA approximates the unbiased estimator. The % variation in Fig. 7A is the normalized eigenvalue from each PC from smartpca, whereas the % variation in 7B represents the proportion of variation explained by population structure. The variances explained by PC1 and PC2 are high and add up to 100% in both A and B since in both cases, since only 2 PCs were utilized.

**Figure 7:**
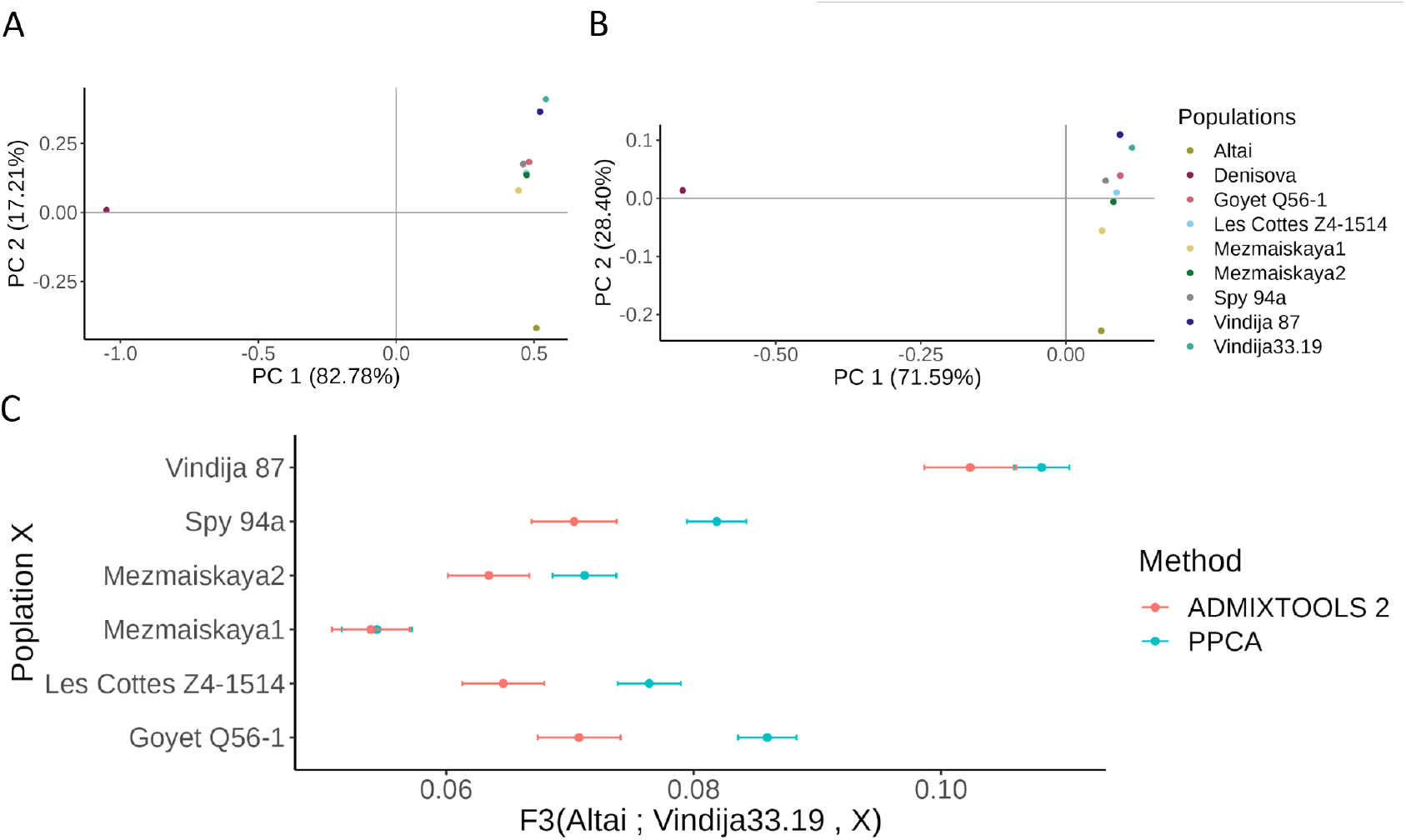
**A.** PCA of archaic neandertals created with smartpca with high coverage neandertals, and projection of the rest. We use two PCs because we just have three high-coverage samples. **B**. PPCA of archaic specimens with scale=2. In A and B, The % variance on x and y-axes denotes the % variance explained by the population structure. **C**. *F*_3_(Altai, Vindija33.19, x) estimated with ADMIXTOOLS 2 and PPCA-based-framework. Larger value on x-axis represents more proximity to Vindija33.19. Bars show 2 standard errors.

We next compare how *f*-statistics computed using PPCA and ADMIXTOOLS 2 differ from each other (Fig 7C). For this purpose, we focus on an outgroup-*F*_3_ statistic *F*_3_(Altai; Vindija33.19, X). This statistic measures the similarity of Neandertal X to the high-coverage Vindija Neandertal, using the Altai Neandertal as an outgroup. More precisely, we project the low-coverage samples onto the line joining Vindija33.19 and Altai in Fig. 7B, and measure the distance from the low-coverage sample to the Altai Neandertal; thus higher values of *f*_3_ denote higher similarity of X to Vindija33.19 (Fig. 2).

Overall, we find the resulting pattern to be similar between the methods, but the PPCA-based values are consistently larger than those obtained using ADMIXTOOLS 2. To further investigate this discrepancy, we focus on Vindia 87, which has been shown to come from the same individual as Vindija 33.19 (Hajdinjak et al., 2018). We decompose *F*_3_ as a linear combination of *F*_2_s:

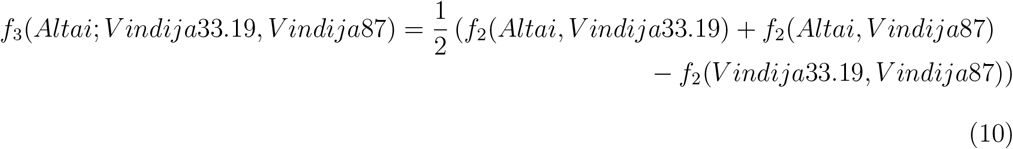

and analyze them seperately for ADMIXTOOLS2 and PPCA. Using ADMIXTOOLS 2, we found that *f*_2_(Altai, Vindija33.19) and *f*_2_(Altai, Vindija 87) have values 0.072 and 0.135, respectively. Since both Vindija samples are from the same individual, the *f*_2_-values are expected to be the identical. In contrast, using the PPCA framework, we obtain the values of *f*_2_(Altai, Vindija33.19)=0.102 and *f*_2_(Altai, Vindija 87)=0.115, respectively, which are much closer. In addition, *f*_2_ should be 0 for two samples from the same individual (i.e. Vindija33.19, Vindija 87). ADMIXTOOLS 2 yields a value of 0.0057, and the PPCA-framework results in a smaller value of 0.00094. Further, we show that f2(Altai, Vindija33.19) is closer to f3(Altai; Vindija33.19, Vindija 87) when estimated from PPCA with 8 PCs compared to that estimated from ADMIXTOOLS 2 (Fig. S8).

One explanation for this is that Vindija 87 is pseudohaploid, and the unbiased estimator in ADMIXTOOLS 2 for *F*_2_ is undefined for single pseudohaploid genomes (*n*_1*s*_ = 1 in eq. 3), and so ADMIXTOOLS2 defaults to a biased estimator. In contrast, the PPCA-based estimator is still defined and can be used in this case, although it introduces the assumption that the samples have the same heterozygosity.

## 4 Methods

### 4.1 PPCA implimentation

We implement PPCA using maximum-likelihood approach following Tipping and Bishop (1999a), and modify this algorithm to work with missing data. Our approach to handle missingness is inspired from EMU (Meisner et al., 2021b). We describe our algorithm briefly:

1. Mean center data **X**_*c*_ = **X** *−* μ.
2. Set missing values to 0.
3. Get covariance matrix 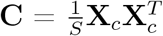
4. Perform eigen value decomposition of **C, C** = **V**Λ**V**^*T*^.
5. Calculate the MLE of the Gaussian noise parameter 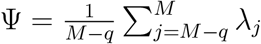 as the average of the *M − q* smallest singular values.
6. Obtain the MLE of the *q*^*th*^ eigenvalue as λ_*q*_ *−* Ψ.
7. Calculate the linear mapping matrix **W** = **V**(Λ *−* Ψ**I**)^1*/*2^.
8. Reconstruct mean-centered data: **X**_**R**_ = **W**(**W**^*T*^ **W**)^*−*1^**W**^*T*^ **X**_*c*_.
9. Replace missing value with reconstructed values.
10. Repeat steps 2-8 until convergence.

Our algorithm differs from that of EMU at steps 4-7, which deal with the modelling of the sampling noise and reconstruction of data, and are specific to PPCA algorithm.

### 4.2 Calculation of standard errors

We use a block-jackknife approach to calculate standard errors (Kunsch, 1989; Maier et al., 2022; Patterson, 2020). We divide the genome in 2 MB blocks, and estimate PCs and *F*-statistics utilizing the entire genotype matrix except for one block. We repeat this procedure for all the blocks. Since the statistics obtained in this way are not independent, we calculate variance similar to ADMIXTOOLS 2 using

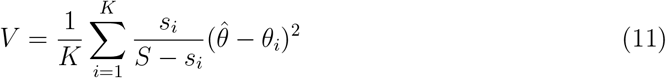

Here, *V* is the variance of a statistic 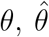 is the average estimate of the statistic, and *θ*_*i*_ is the estimate with block *i* removed. Here *K* denotes the number of blocks, *s*_*i*_ the number of sites in block *i*, and *S* is the total number of sites.

### 4.3 Simulation pipeline

We used slendr (Petr et al., 2022) to simulate 4 populations with 11 individuals each and 6 populations with 10 individuals each. We used a mutation rate of 10^*−*8^ per base per generation, a recombination rate of 10^*−*8^ per base per generation, and a generation time of 30 years. We set the effective population size for each population to 1000. Five of these populations were created by admixing other populations to create a complex scenario. Additionally, we created a geneflow event from *X*_3_ to *X*_2_ 500 generations ago (Fig. 2). We calculated true *F*-statistics from the simulated samples using the branch lengths of the trees, using the functions implemented in `tskit` (Baumdicker et al., 2022). We used eigenstrat files as input for all PCA methods and ADMIXTOOLS 2.

## 5 Discussion

In this study, we present a statistical framework to jointly compute PCA and *F*-statistics. Many ancient genetic studies use both of these tools, but make slightly different assumptions, slightly different models and different software for them. In contrast, our joint framework allows us to make sure that assumptions are consistent throughout the analysis. The key advantage is that the effect of modelling assumptions becomes apparent, and this also allows us to make novel recommendations about how PCA-based analyses should be performed and interpreted.

The connections between *F*-statistics and PCA allow us to provide a better understanding on how different PCA algorithms emphasize different parts of the data, and how they emphasize population structure versus sampling noise.

In particular, *F*-statistics enable quantitative interpretation of PCA-plots, where distances on a PCA are directly proportional to genetic differentiation, and orthogonal projections can directly be used to test for admixture. However, the above statement is true only when all the (relevant) PCs are used. Visualizing two or only few PCs may be insufficient to accurately visualize population structure and admixture.

### 5.0.1 Interpreting PCA plots

There is a considerable literature aimed at interpreting PCA-plots, which generally fall in two approaches: First, for simple models, such as lattice models (Novembre and Stephens, 2008) and isolation-without-migration models (McVean, 2009), the eigenvectors, and hence the PCs, can be calculated analytically. These approaches are powerful to understand the “inner workings” of PCA, but do not deal with the variation and noise inherent in sample data. Thus, much more common are interpretations based on empirical observations, such as that the first PC commonly aligns with the axis of an expansion (Cavalli-sforza et al., 1996), that PCs tend to align with the branches of a population tree or that the first two PCs recap a map of Europe (Novembre et al., 2008; Cavalli-Sforza and Piazza, 1975).

However, these observational guidelines have provided to not hold particularly well in simulations, and have been shown to be strongly influenced by sampling schemes, simulation details and other factors (Novembre and Stephens, 2008; DeGiorgio and Rosenberg, 2013; Jay et al., 2013) enough that interpretations of the first few PCs in terms of demographic history are frequently discouraged Elhaik (2022).

Our link between PCA and *F*-statistics provides a different way of interpreting PCA-plot, by studying PCA in terms of the embedded *F*-statistics. A key difference is that our approach is quantitative, and provides exact results when all PCs are used, and good approximations when only the first few PCs are used (Peter, 2022). Then, *F*-statistics can directly be calculated from one-dimensional PCA-plots such as that in Figure 3B: *F*_2_(*X*_1_, *X*_2_) corresponds to the (squared) sum of the distance between *X*_1_ and *X*_2_ in that figure, and we can see from the plot that PC6 is the main PC that teases out that axis of variation (because the two populations are furthest apart). The drawback of our approach is that *F*-statistics are not directly interpretable in terms of demographic history, and we need additional steps to link them with what we are typically interested in.

### 5.0.2 Comparing PCA methods

Different PCA methods we discussed here differ in how they deal with biological vs. sampling variation, and how different sources of variations are emphasized. This gives us some insights for when different PCA algorithms should be used.

Classical PCA is mathematically the simplest, and still the most widely used method for visualizing population structure. It has an interpretation in terms of pairwise coalescent times (McVean, 2009). However, because classical PCA incorporates *all* variation in the data, it does not distinguish between variation due to population structure, and variation due to sampling. This can lead to problems: For example, one cannot directly project additional samples to a PCA, instead some correction is required otherwise some “shrinkage” occurs where new samples are projected closer to the origin (Patterson et al., 2006; Wang et al., 2015). Thus, in our opinion classical PCA should be the primary choice for analyses primarily aimed at quality control, because incorporating noise allows it to reveal technical artifacts and outliers. Effects of different sequencing or capturing techniques will very often be visible on a PCA-plot, while they may be more hidden and only partially corrected in PPCA.

Our results suggest that the methods that separate sampling noise from population structure, such as PPCA and LSE, are preferable to classical PCA when the primary goal is to depict population structure. PPCA is the simpler of the two methods, because it assumes homoskedastic noise, i.e. that all samples have the same heterozygosity or sampling variance. As Tipping and Bishop (1999a) showed, without missing data, the maximum-likelihood estimator of PPCA results in virtually identical PCA plots compared to those obtained from a classical PCA; the only differences are that the axes for the first few PCs will be re-scaled, and that the majority of PCs are set to zero (Fig. S1, Fig. S2).

LSE goes one step further by modelling heteroskedastic, binomial noise. This ensures that both PCA and *F*-statistics use the exact same modelling assumptions, and thus LSE-based PCA plots are directly comparable to *F*-statistics (Fig. S4). This relationship is exact if all PCs are used. One advantage of LSE is that the eigenvalues corresponding to the PCs irrelevant for population variation have an expectation of zero. Since these eigenvalues are directly proportional to the amount of variance explained per PC, and to the contribution to *F*-statistics, we expect that truncating them will yield good results, which is indeed what we see in simulation results (Fig. S3).

In real data, this expectation of zero for the eigenvalues will typically yield both positive and negative eigenvalues, which is why the equation for computing *f*_2_ needs to be adjusted (eq. 9), and subtract the contribution to *F*_2_ by PCs that have negative eigenvalues.

From a theoretical point of view, the model optimized in LSE is more desirable to that in PPCA, because it takes into account that different individuals might have different heterozygosities. However, the advantage of PPCA is that it is relatively easier to implement with missing data, even for single pseudohaploid samples, which are common in ancient DNA applications (Fig. 5) (Tipping and Bishop, 1999a; Orlando et al., 2021).

A further approach that is used for ancient DNA is to project low-coverage data onto a “reference”-data set. This has three useful advantages: First, the overall shape of the PCA only depends on the reference data set. Thus, PCA-plots using the same reference data become directly comparable, which can be useful as a quick way of assessing population variation, e.g. using the Western Eurasian PCA used in studies of Holocene ancient DNA studies from Europe and Western Asia (Haak et al., 2015). Second, it deals with missing data in the projected samples, because often only a subset of sites are required for an accurate projection. Third, it also deals with sampling noise in the projected sample, because the sampling noise is orthogonal to the variation in the reference data set, and thus gets removed by the projection.

However, the drawback of using projections, and why they, in our opinion, are inferior to probabilistic PCA, is that they do not capture the full variation in the data. In particular, only the variation in the reference data is considered, but not the variation that is private to the projected samples.

In terms of *F*-statistics (Peter, 2022),

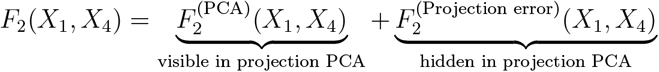

This is particularly problematic when projecting ancient human samples onto a modern reference data set, because the differentiation between ancient human population is often considerably larger than that between present-day populations (Haak et al., 2015; Lazaridis et al., 2014), and thus projection PCA have no quantitative interpretation.

### 5.0.3 Plotting PCs

Our connections between *F*-statistics and PCA in a joint framework directly suggest some recommendations for how PCs should be plotted when displaying population genetic variation.

First, the scale at which PCs are plotted matters, and the units on the different PCs should be comparable (Figure 3B). Thus, we recommend that PCA-biplots should be plotted with “fixed” aspect ratios where the x- and y-axes have the same units and scale, contrary to the current common practice where the axes are scaled arbitrarily (e.g. Novembre et al. (2008); Peter et al. (2020)). It also highlights the benefits of plotting PCs on a shared axis such as in Figure 3B, where the decreased variance explained by each subsequent PC is directly visualized, and tens of PCs can be plotted jointly.

Second, it has been a “best-practices” suggestion that PCs are plotted along with the proportion of variance explained by each PC (e.g.(Novembre and Peter, 2016; Elhaik, 2022)). While we think this holds, it is important to recognize that different PCA algorithms handle this differently: In classical PCA, the proportion of variance is out of the total variance, including population structure, within-population variation and sampling noise. In contrast, in PPCA the number of PCs retained is a parameter, which *sets* the proportion of variance explained by population structure. Thus, we obtain the (user-controlled) proportion of variance due to noise (corresponding to Ψ), and the proportion of variance each PC explains as a proportion of the variation explained by population structure. For LSE, the proportion of variance is calculated from the variance contributing to the population structure. In addition, the proportion of variance explained from a projected PCA will depend on the number of PCs or eigenvectors used for reconstruction of projected data (Patterson et al., 2006). Thus, the percent of variation explained between classical PCA, PPCA, LSE and projected PCA are not directly comparable.

### 5.0.4 Grouping individuals

A benefit of the interpretation that we developed is that it demonstrates when and how we can use PCA to group individuals into populations for *F*-statistics-based analysis. Thus, the groupings of individuals into populations becomes a result of the analysis, as opposed to an *a priori* assumption set by the researcher.

This *a priori* grouping may lead to difficulties in analyses, for example when dealing with recent migrants, or when grouping sparsely sampled populations (Shringarpure and Xing, 2014). As ancient genetic sampling becomes more dense, recent migrants have become increasingly prevalent in data sets. These migrants are important, because they allow tracking of movements – often over large distances – within the last handful of generations. Thus, these recent migrants have become an important focus of study and it is desirable to distinguish recent migration from the more distant-in-time migration that is ordinarily the focus of *F*-statistic analyses.

A PCA will reveal such recent migrants as outliers of that group with their population-of-origin, whereas they would be missed by *F*-statistics.

In sparsely sampled populations, it is often necessary to group individuals that span relatively large areas or times to get to a usable sample size, particularly for regions where DNA preservation is poor. However, grouping genetically distinct individuals can be risky, as population substructure can invalidate the interpretation of *F*-statistic-tests (Peter, 2016). However, PCA provides a meaningful and simple way to assure that populations are devoid of relevant substructure: If individuals cluster tightly on *all* PCs that are meaningful for between-cluster variation, they can safely be grouped. However, if there is within-population variation that is correlated with other populations, then *F*-statistic-tests cannot be used to test for admixture.

As discussed above, these considerations hold particularly when using a PCA method that separates sampling noise from population structure, such PPCA or LSE. For classical PCA, clustering will also be impacted by sample quality and depth - which can be separated out effectively by the probabilistic methods.

### 5.0.5 Handling missing data

Incomplete genotype calls with missing data are common in ancient DNA, and thus it is important that methods are able to deal with it. Classical PCA, which is based on SVD, relies on complete data, although methods that impute data are becoming more common (Meisner et al., 2021b). For *F*-statistics, ADMIXTOOLS 2 has a parameter `maxmiss` that controls how much missingness is permitted, e.g. `maxmiss=0` removes all sites with any missing data, and `maxmiss=1` retains all the sites with any data.

For our tests with missingness, we could not use the option maxmiss=0, since this resulted in no sites available. Hence, we used `maxmiss=1` throughout, which resulted in inflated *F*-statistics in our simulations with 50% missing genotype calls. In contrast, the PPCA-based algorithm is not affected, and appears to be more robust to missingness for both 8 and 12 PCs. It may seem surprising that ADMIXTOOLS 2 shows inflated estimates of *F*-statistics, given that it is based on Patterson’s unbiased estimator. The reason for this inflation is that in the analysis for missing data we use one individual from each population, and when there is missing data, it results in only haploid genotype. In such cases, the correction term for the unbiased estimator is undefined. Thus we show that, PPCA provides a practical solution in case of estimation of individual-based *F*-statistics with missing data.

### 5.0.6 Application on real data

We used a PPCA-based framework with Neandertal data to estimate PCs utilizing all the samples. This approach is more straight-forward than the authors’ approach to first estimate the PCs using high-coverage genomes, and then project the low-coverage genomes. We show that one can accurately estimate outgroup *F*_3_ statistic from the PCs. We find that *F*_2_ calculated using diploid genotype data with our framework is comparable to that of ADMIXTOOLS 2. However, ADMIXTOOLS 2 can not be used to estimate *F*_2_ with pseudohaploid data since the unbiased estimator in ADMIXTOOLS 2 is undefined in this case. We show that PPCA-based framework provides accurate estimates even with pseudohaploid samples. Here, we point out that the reason PPCA performs well in this case is because the scalar noise parameter ψ is determined from all the samples.

### 5.0.7 Limitations and future directions

One limitation of this framework is the need to perform multiple PPCAs to obtain standard errors, which is computationally expensive. Further studies are needed to design a statistical framework that can estimate the errors using SNP loadings, and therefore can work fast with large datasets. In case of PPCA, one needs to determine the number of PCs relevant for the particular analysis. We show that PPCA is not sensitive to small changes in the number of PCs, and we provide the option of LSE which does not require user-defined number of PCs. Finally, for LSE, an algotithm to deal with missing data is still pending.

### 5.0.8 Summary

To summarize, we present a method to perform PCA and *F*-statistics jointly and show that this approach not only improves estimates of *F*-statistics, but also provides a solution to the standardization and quantification of PCA. Our framework is available on github as a snakemake pipeline Popli (2024).

## 6 Appendix

Here, we show that LSE Cabreros and Storey (2019) leads to unbiased *F*-statistics. Following Cabreros and Storey (2019), we define a symmetric matrix **G**:

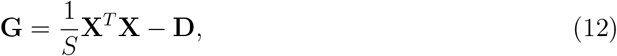

Here, **X** is the (uncentered) allele frequency matrix with shape *S × M*, **D** is a matrix of diagonal entries *d*_*ii*_ “correcting” heterozygosities, M is the number of individuals and *S* is the number of SNPs.

In particular *d*_*ij*_ = 0 for *i≠ j* and,

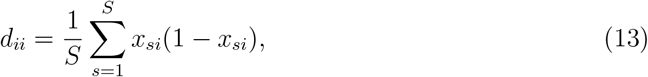

for diploid data. This is same as the correction term given by Reich et al. (2009) and Patterson et al. (2012), except here we are calculating individual-based covariance matrix, and so the denominator term *n*_*si*_ *−* 1 = 1, where *n*_*si*_ is the number of haploids.

Thus,

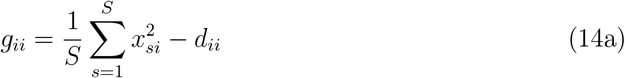

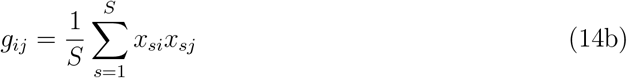

Consider now

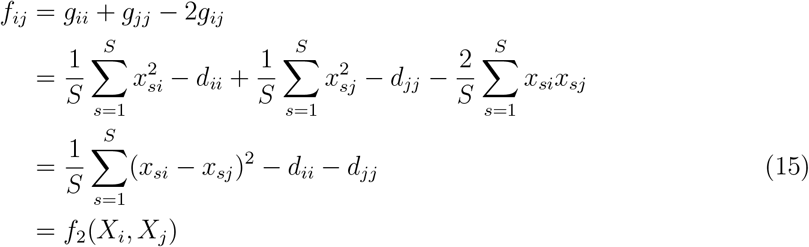

Hence the matrix **G** can be used to estimate unbiased *F*_2_-statistics.

### 6.1 Availability of data and materials

An open-source implementation of our snakemake pipeline to estimate PCA and *F*-statistics, along with a toy example dataset, and the scripts to generate our simulations are available on GitHub Popli (2024). The analyzed Neandertal dataset was generated in previous studies, and is available in European Nucleotide Archive Hajdinjak et al..

